## Supplemental Material for "Systematic tissue annotations of –omics samples by modeling unstructured metadata"


| | Page |
|---|---|
| Supplemental Figures | 2 |
| Supplemental Tables | 12 |
| Supplemental Notes | 15 |

### Supplemental Figures

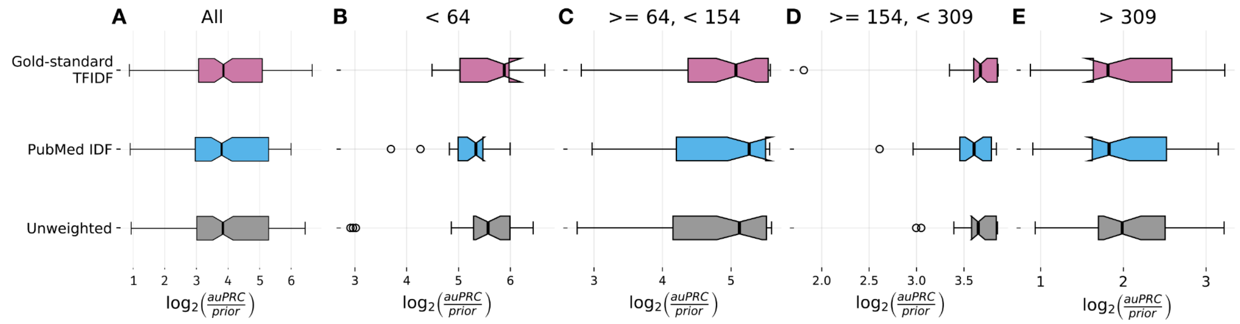


**Figure S1. NLP-ML models trained using sample embeddings created using three different word-weighting schemes have similar performance in sample tissue classification.** The three word-weighting schemes are along the y-axis. Gold-standard TFIDF: weights are based on term-frequency inverse-document-frequency (TFIDF) values calculated from descriptions for samples in our gold standard. PubMed IDF: weights are based on IDF values calculated from the entirety of PubMed. Unweighted: weights are all equal to 1. Performance is shown on the x-axis as the logarithm of the area under the precision-recall curve (auPRC) over the prior, where the prior is the fraction of positive over positive and negative training examples. This metric accounts for the variable number of annotated terms per tissue. Each boxplot shows the distribution of this metric across tissues. Panel A shows the results for all tissue and panels B–E show the same results broken down by number of training examples per tissue (indicated at the top of each plot).

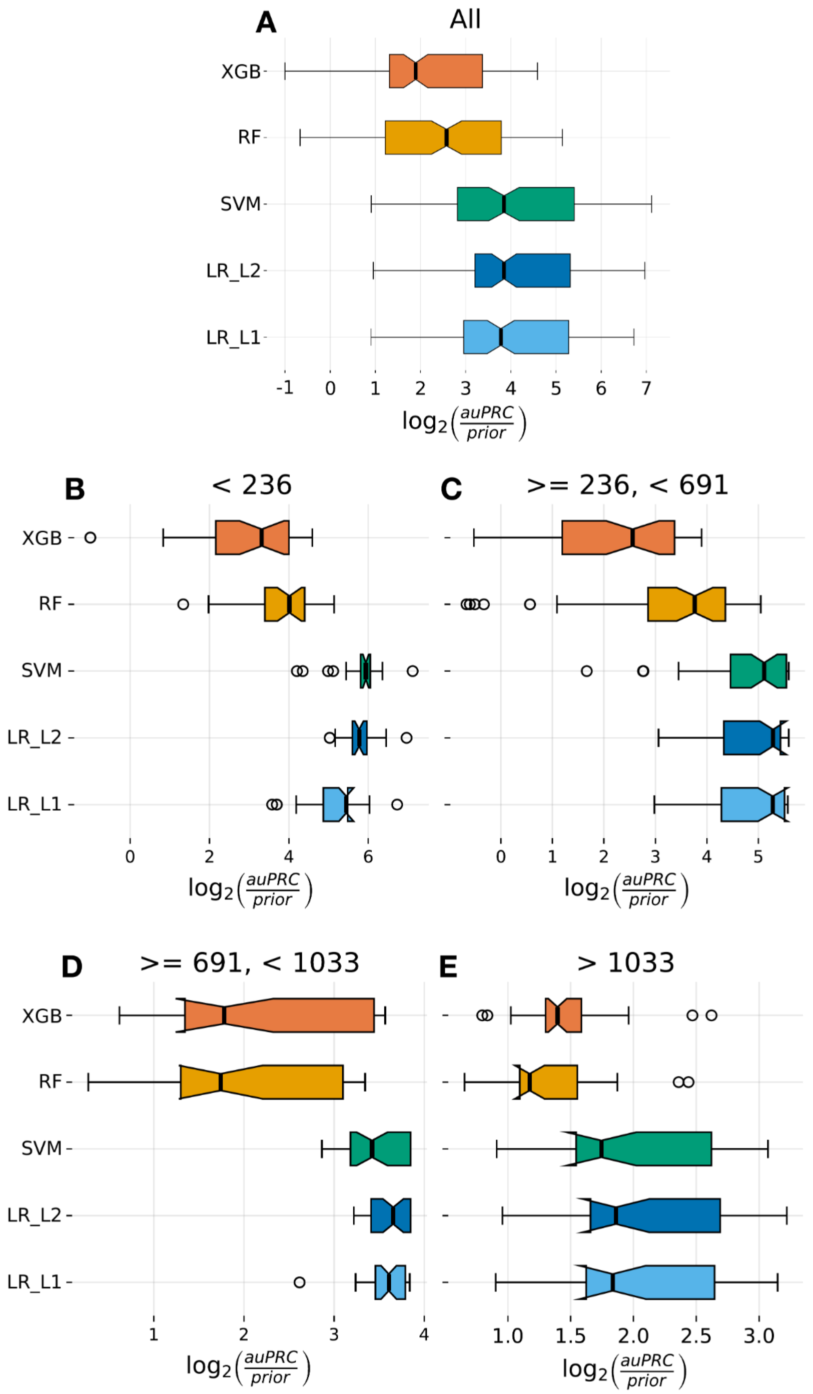


**Figure S2. Logistic regression provides the best performance among different machine learning classifiers for building NLP-ML models.** The five classifiers are along the y-axis. XGB: XGBoost. RF: Random Forest. SVM: Support Vector Machine. LR_L1/L2: Logistic Regression with L1 or L2 regularization. Performance is shown on the x-axis as the logarithm of the area under the precision-recall curve (auPRC) over the prior (where the prior is the fraction of positive over positive and negative training examples) calculated as the average over 3-fold cross validation. Each boxplot shows the distribution of this metric across tissues. Panel A shows the results for all tissue and panels B–E show the same results broken down by number of training examples per tissue (indicated at the top of each plot).

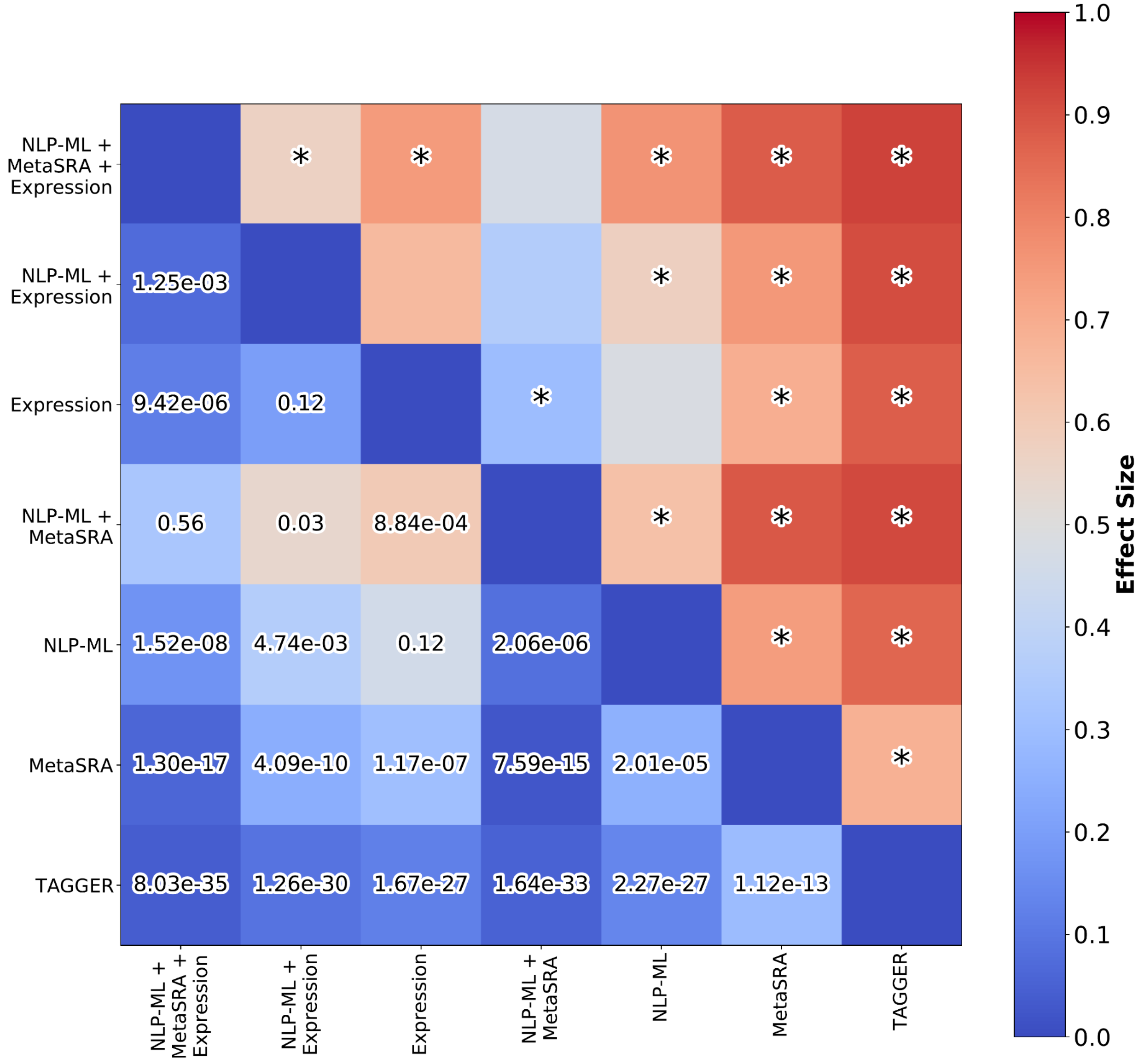


**Figure S3. NLP-ML and its combination with other methods provide substantial and statistically significant improvements in sample tissue annotation.** The heatmap shows the results of comparison between all pairs of methods. The color of each cell shows the proportion of tissues where the method along the row outperformed the method along the column. An asterisk in the cell for the upper-right triangle of the matrix denotes a statistically significant difference between the row and column methods (corrected $p < 0.01$) determined using a Wilcoxon rank sum test with Benjamini-Hochberg correction for multiple hypothesis testing. The actual p-values are shown in the lower-left triangle. Methods are ordered based on the number of other methods they differ significantly from.

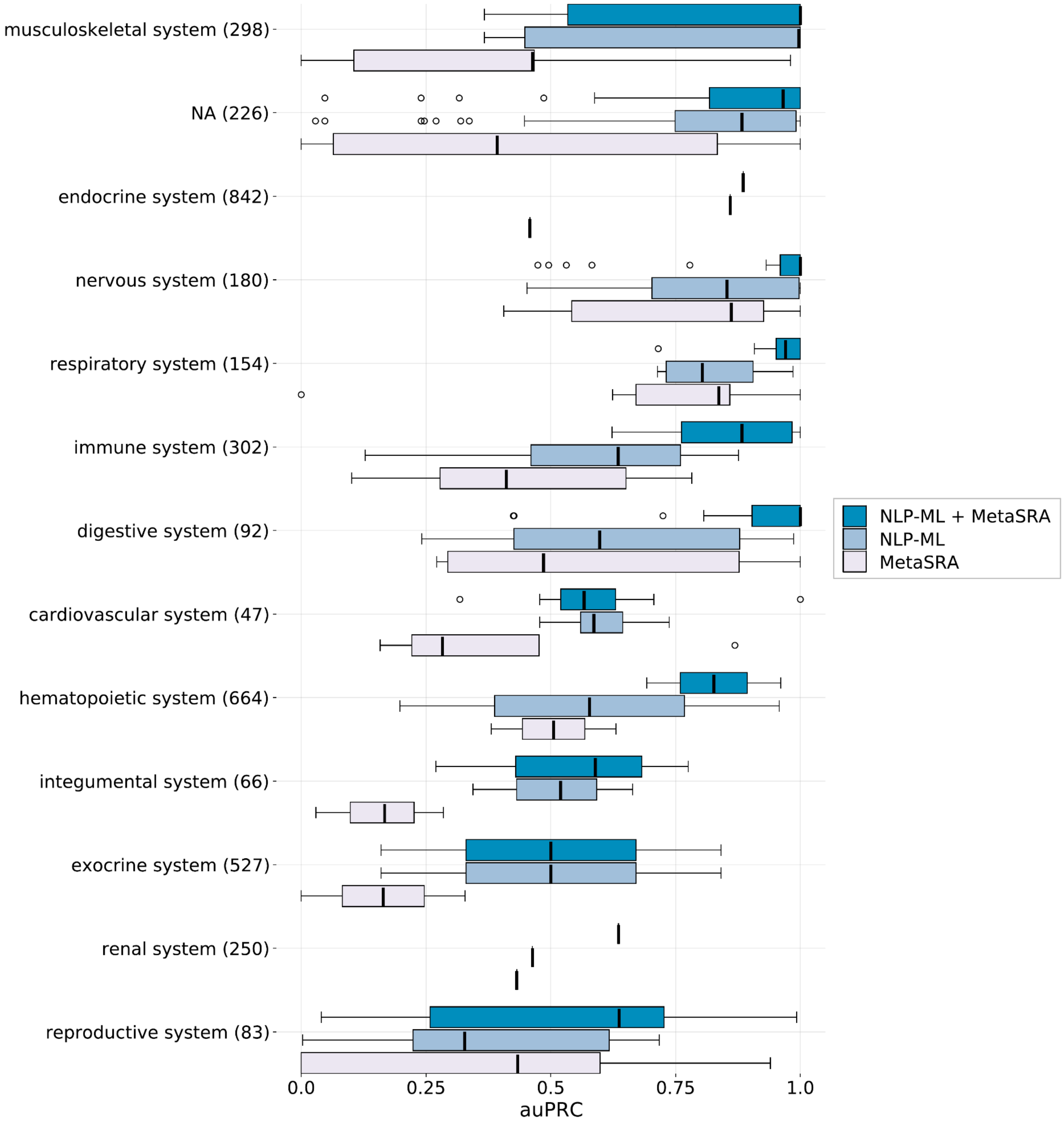


**Figure S4. Relative performance of NLP-ML and MetaSRA are nearly consistent across tissues in various high-level anatomical systems.** Each boxplot is the distribution of the area under the precision-recall curve (auPRC) scores across tissues for MetaSRA, NLP-ML, and their combination ('NLP-ML+MetaSRA'). Each point in the boxplot is the performance for a single tissue model averaged across cross validation folds. Each group of boxplots corresponds to the set of tissues pertaining to a particular high-level anatomical system (y-axis).

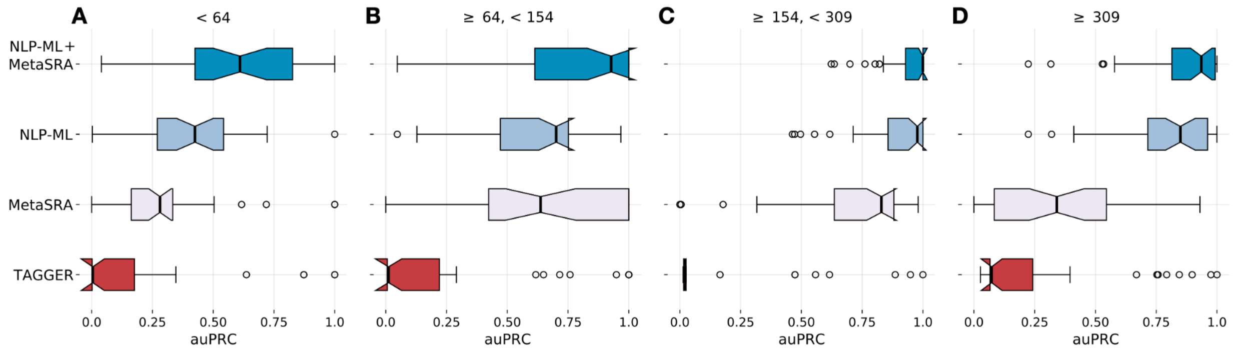


**Figure S5. The performance of NLP-ML over and above other text-based methods varies across tissues with different training set sizes.** Distribution of the area under the precision-recall curve (auPRC) scores across tissues for each of the three individual text-based methods for sample classification: TAGGER, MetaSRA, and NLP-ML. Also shown is the distribution of auPRC scores for combining the predictions of NLP-ML and MetaSRA. The panels show the same results in **Figure 2** broken down by number of training examples per tissue (indicated at the top of each plot).

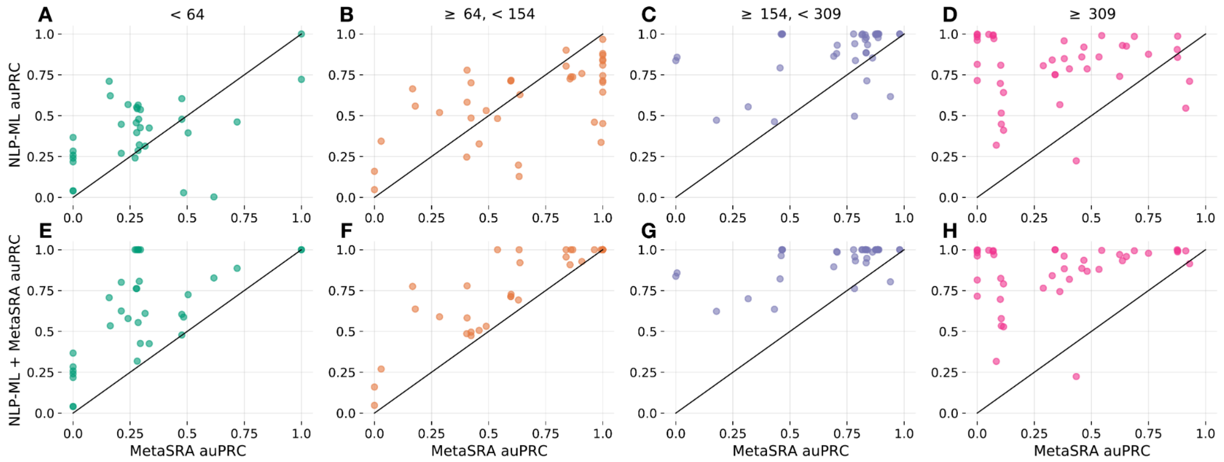


**Figure S6. Despite variability in the relative performance of NLP-ML and MetaSRA across tissues with different training set sizes, combining them consistently results in good performance.** Each scatterplot shows the area under the precision-recall curve (auPRC) scores of sample tissue predictions by two methods labelled on the x- and y-axis. The panels show the comparison for tissues grouped by number of training examples per tissue (indicated at the top of each plot). A–D) Comparison of MetaSRA (x-axis) vs. NLP-ML (y-axis). E–H) Comparison of MetaSRA (x-axis) vs. the combination of predictions from NLP-ML and MetaSRA.

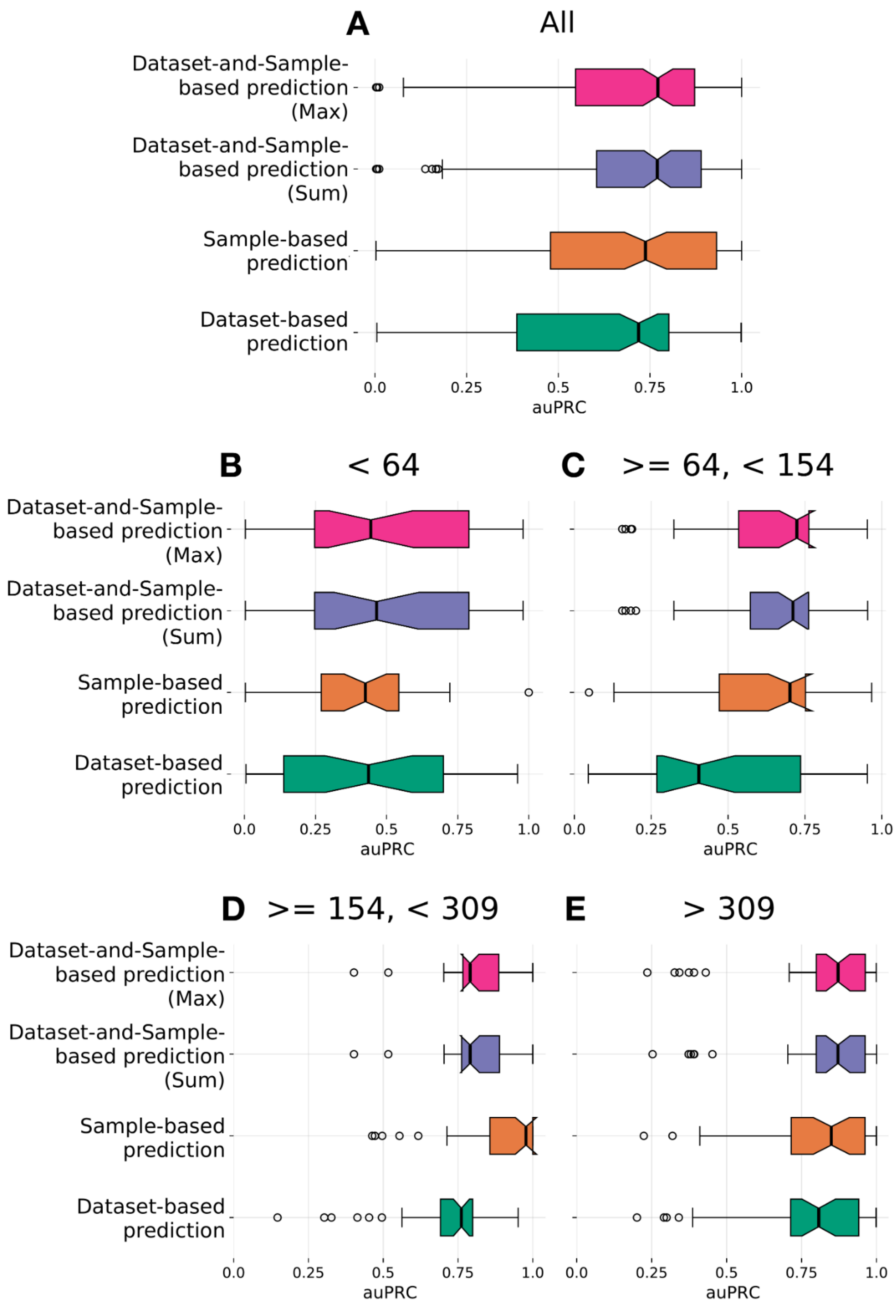


**Figure S7. The relative performance of NLP-ML predictions based on sample description, dataset description, or their combination varies by training set size.** The description sources are along the y-axis. The combination of predictions from the dataset and the sample descriptions were done either by summing the predictions (Sum) or by taking the maximum among the two (Max). Performance is shown on the x-axis as the area under the precision-recall curve (auPRC). Each boxplot shows the distribution of this metric across tissues. Panel A shows the results for all tissue and panels B–E show the same results broken down by number of training examples per tissue (indicated at the top of each plot).

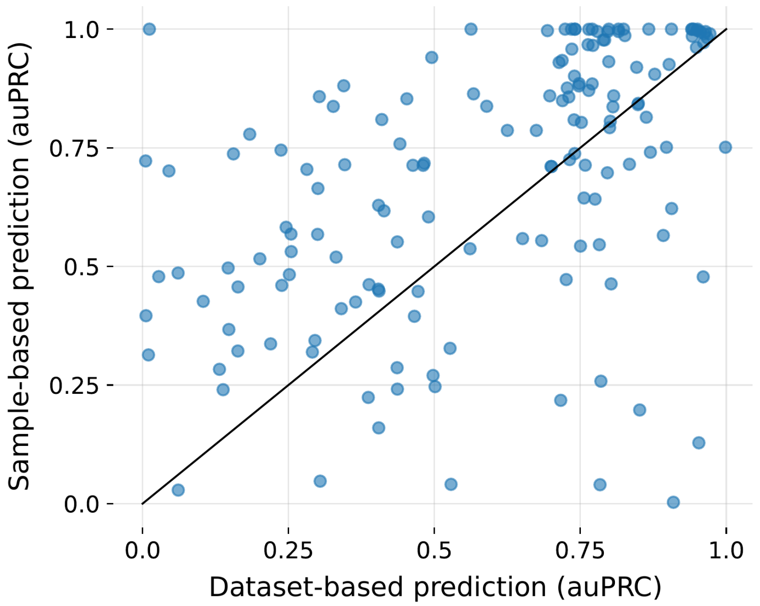


**Figure S8. NLP-ML sample annotations based on sample description invariably outperforms sample annotations based on dataset description.** Scatterplot of the area under the precision-recall curve (auPRC) scores of sample tissue predictions from sample text (x-axis) vs. predictions from dataset text (y-axis). The dataset-based prediction is made by assigning the predicted probability for the dataset description to all samples in that dataset. Each point in the scatterplot correspond to a tissue/cell-type term. The solid line denotes equal performance between the two methods.

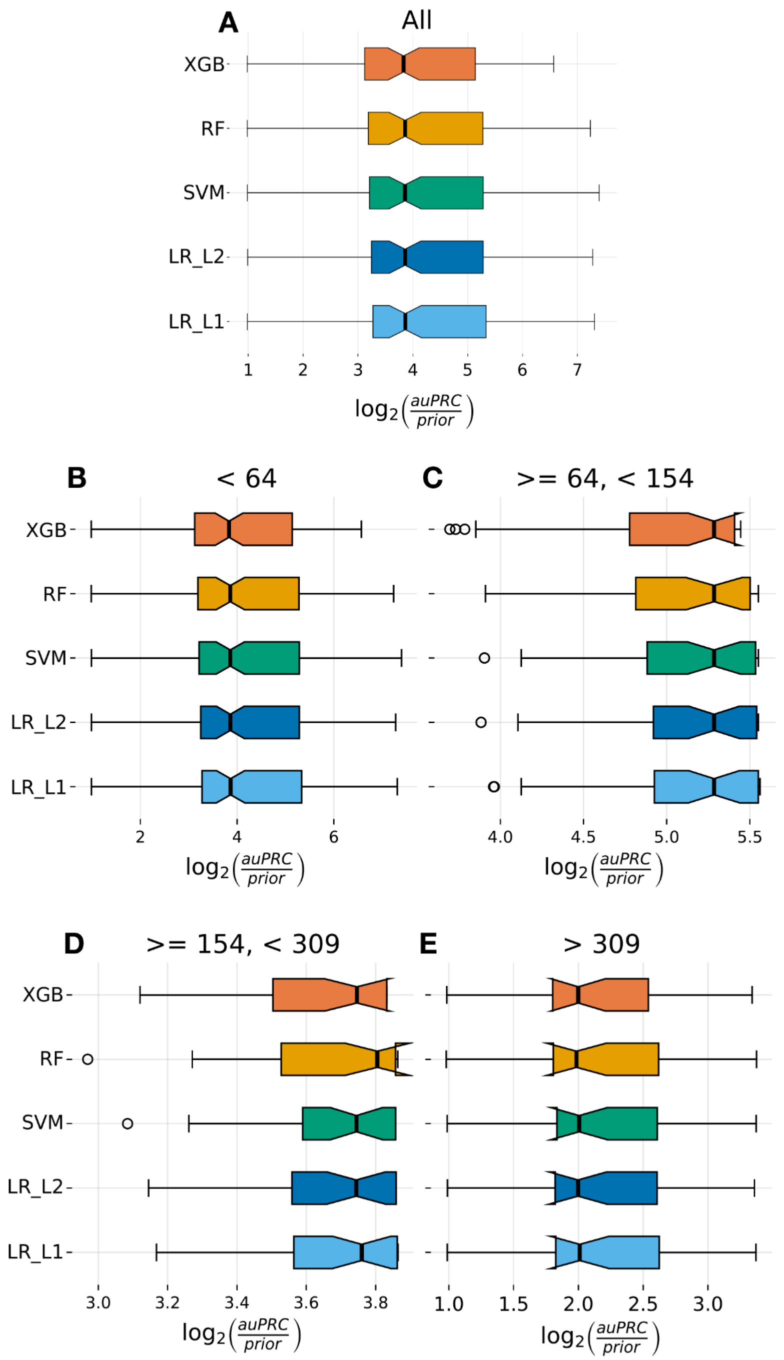


**Figure S9. Logistic regression provides slightly better performance among different machine learning classifiers for building expression-based tissue annotation models.** The five classifiers are along the y-axis. XGB: XGBoost. RF: Random Forest. SVM: Support Vector Machine. LR_L1/L2: Logistic Regression with L1 or L2 regularization. Performance is shown on the x-axis as the logarithm of the area under the precision-recall curve (auPRC) over the prior (where the prior is the fraction of positive over positive and negative training examples) calculated as the average over 3-fold cross validation. Each boxplot shows the distribution of this metric across tissues. Panel A shows the results for all tissue and panels B–E show the same results broken down by number of training examples per tissue (indicated at the top of each plot).

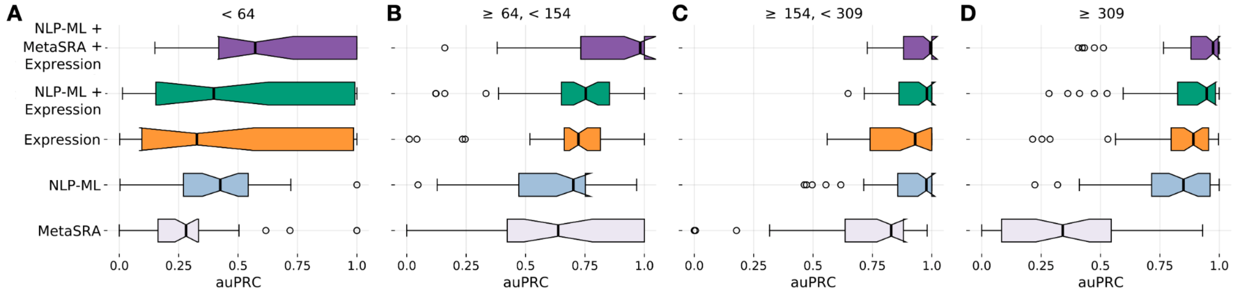


**Figure S10. Relative performance of text-based and expression-based methods varies across tissues with different training set sizes.** Distribution of the area under the precision-recall curve (auPRC) scores across tissues for the two top-performing text-based methods – MetaSRA and NLP-ML – and for the method based on expression profiles ('Expression') for sample tissue classification. Also shown are the distributions of auPRC scores for combining the predictions of Expression with NLP-ML ('NLP-ML+Expression') and with NLP-ML and MetaSRA ('NLP-ML+MetaSRA+Expression'). Each point in the boxplot is the performance for a single tissue model averaged across cross validation folds. The panels show the same results in **Figure 6A** broken down by number of training examples per tissue (indicated at the top of each plot).

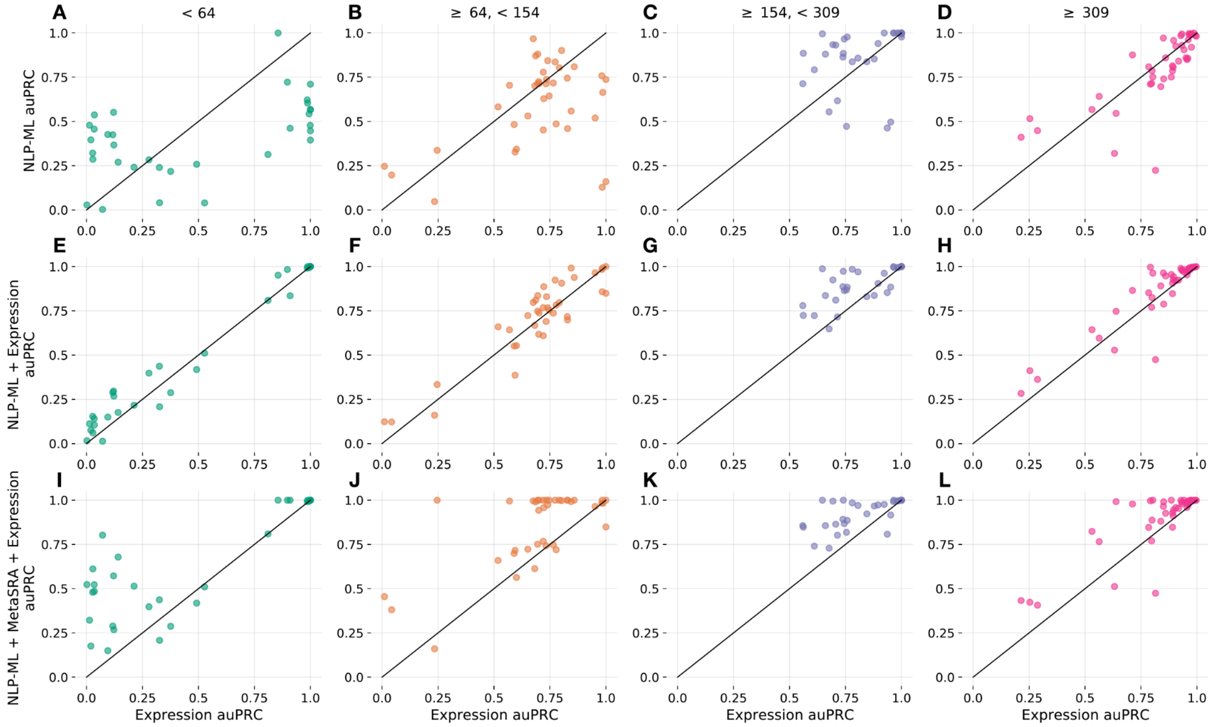


**Figure S11. Despite variability in the relative performance across tissues with different training set sizes, combining text-based and expression-based models results in consistently good performance.** Each scatterplot shows the area under the precision-recall curve (auPRC) scores of sample tissue predictions by expression-based models on the x-axis vs another model/method on the y-axis. The panels show the comparison for tissues grouped by number of training examples per tissue (indicated at the top of each plot). A–D) Comparison to NLP-ML (y-axis). E–H) Comparison to the combination of predictions from NLP-ML and Expression. I–L) Comparison to the combination of NLP-ML, MetaSRA, and Expression.

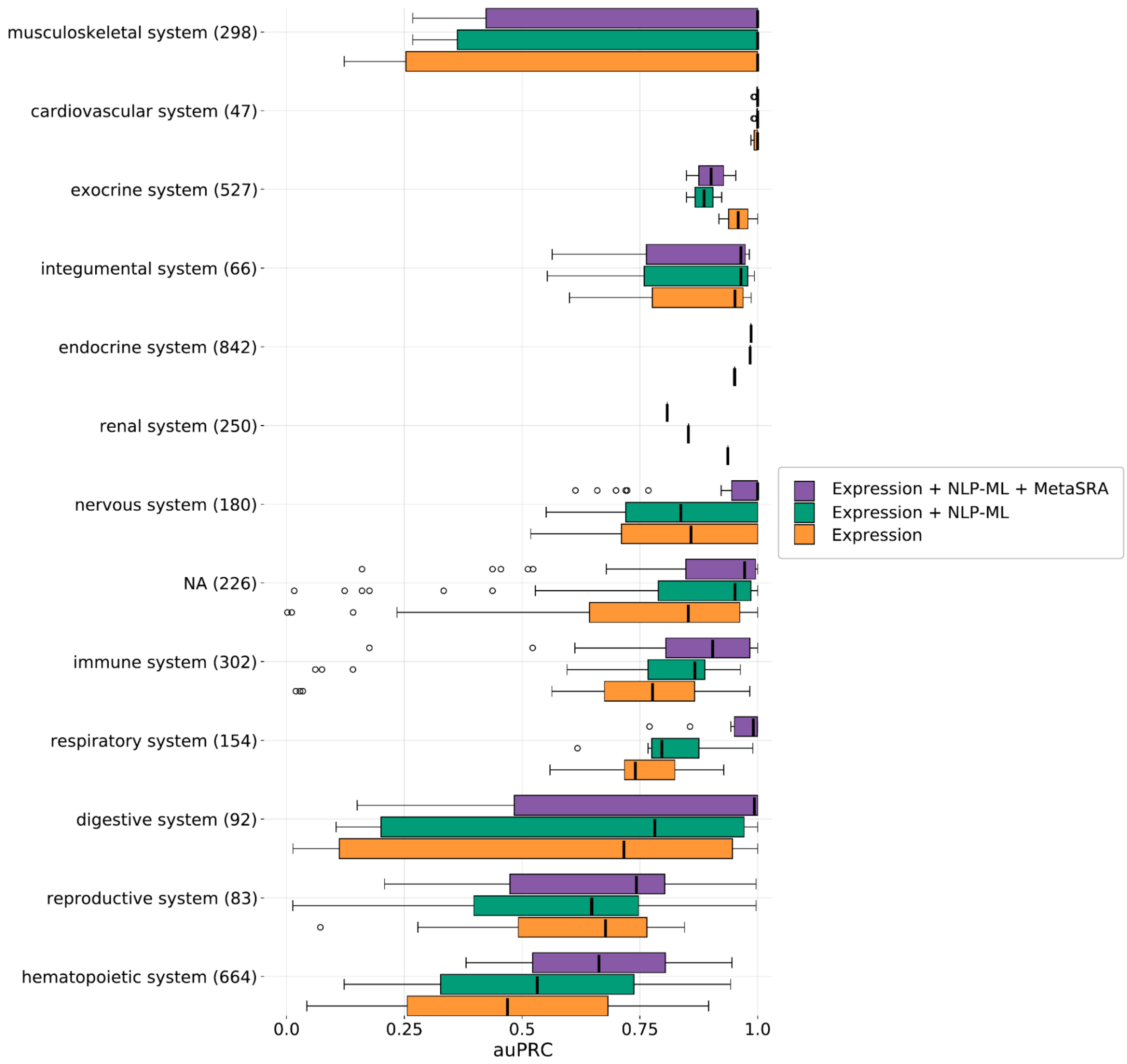


**Figure S12. Relative performance of text-based and expression-based models are nearly consistent across tissues in various high-level anatomical systems.** Each boxplot is the distribution of the area under the precision-recall curve (auPRC) scores across tissues for Expression, MetaSRA, NLP-ML, and their combinations ('Expression+NLP-ML' and 'Expression+NLP-ML+MetaSRA'). Each point in the boxplot is the performance for a single tissue model averaged across cross validation folds. Each group of boxplots corresponds to the set of tissues pertaining to a particular high-level anatomical system (y-axis).

#### Supplemental Tables

**Table S1.** Predictions from liver (UBERON:0002107) model on list of GOBP terms relevant to the liver.

| GO biological process term | Probability |
|---|---|
| hepatic immune response | 0.99 |
| liver regeneration | 0.99 |
| positive regulation of hepatocyte proliferation | 0.99 |
| hepatocyte apoptotic process | 0.99 |
| positive regulation of hepatic stellate cell activation | 0.99 |
| regulation of hepatocyte growth factor receptor signaling pathway | 0.99 |
| positive regulation of hepatocyte growth factor receptor signaling pathway | 0.99 |
| negative regulation of hepatocyte growth factor biosynthetic process | 0.99 |

**Table S2.** Predictions from skeletal muscle tissue (UBERON:0001134) model on list of GOBP terms relevant to the skeletal muscle tissue.

| GO biological process term | Probability |
|---|---|
| musculoskeletal movement | 1.00 |
| regulation of skeletal muscle contraction by action potential | 1.00 |
| voluntary musculoskeletal movement | 1.00 |
| regulation of skeletal muscle contraction | 1.00 |
| skeletal muscle tissue growth | 1.00 |
| regulation of skeletal muscle satellite cell proliferation | 1.00 |
| regulation of slow-twitch skeletal muscle fiber contraction | 1.00 |
| regulation of skeletal muscle cell differentiation | 1.00 |
| myofibril assembly | 1.00 |
| regulation of skeletal muscle tissue development | 0.99 |
| positive regulation of fast-twitch skeletal muscle fiber contraction | 0.99 |
| skeletal muscle myosin thick filament assembly | 0.99 |
| regulation of striated muscle contraction | 0.99 |
| negative regulation of skeletal muscle cell differentiation | 0.98 |
| regulation of the force of skeletal muscle contraction | 0.98 |
| regulation of skeletal muscle contraction via regulation of action potential | 0.97 |
| skeletal muscle thin filament assembly | 0.97 |
| skeletal muscle satellite cell differentiation | 0.97 |

| | |
|---|---|
| negative regulation of skeletal muscle tissue growth | 0.96 |
| positive regulation of skeletal muscle cell differentiation | 0.96 |
| negative regulation of skeletal muscle tissue development | 0.94 |
| positive regulation of skeletal muscle tissue development | 0.94 |
| diaphragm development | 0.92 |
| negative regulation of striated muscle contraction | 0.90 |
| striated muscle atrophy | 0.86 |
| regulation of muscle contraction | 0.81 |
| relaxation of skeletal muscle | 0.74 |
| skeletal muscle cell differentiation | 0.49 |
| regulation of smooth muscle cell proliferation | 0.49 |
| negative regulation of striated muscle tissue development | 0.48 |
| regulation of skeletal muscle adaptation | 0.41 |
| growth factor dependent regulation of skeletal muscle satellite cell proliferation | 0.39 |
| negative regulation of myoblast differentiation | 0.38 |
| myoblast proliferation | 0.32 |
| extraocular skeletal muscle development | 0.31 |
| positive regulation of myoblast differentiation | 0.25 |
| regulation of skeletal muscle contraction by regulation of release of sequestered calcium ion | 0.25 |
| positive regulation of striated muscle tissue development | 0.20 |
| skeletal muscle tissue regeneration | 0.19 |
| positive regulation of skeletal muscle fiber development | 0.19 |
| myoblast migration | 0.19 |
| diaphragm contraction | 0.15 |
| myoblast fate commitment | 0.13 |
| positive regulation of skeletal muscle contraction by regulation of release of sequestered calcium ion | 0.11 |
| striated muscle contraction | 0.10 |
| skeletal muscle contraction | 0.09 |
| myoblast fusion | 0.08 |
| myoblast differentiation | 0.06 |
| skeletal muscle fiber development | 0.05 |

**Table S3.** Predictions from brain (UBERON:0000955) model on list of DO terms relevant to the brain.

| DO disease term | Probability |
|---|---|
| neuronal ceroid lipofuscinosis | 1.00 |
| complex cortical dysplasia with other brain malformations | 1.00 |
| neurodegeneration with brain iron accumulation | 1.00 |
| hypomyelinating leukodystrophy | 1.00 |
| Parkinson's disease | 1.00 |
| Joubert syndrome | 1.00 |
| Ritscher-Schinzel syndrome | 1.00 |
| holoprosencephaly | 1.00 |
| autosomal dominant nocturnal frontal lobe epilepsy | 0.99 |
| advanced sleep phase syndrome | 0.97 |
| Warburg micro syndrome | 0.67 |
| Meckel syndrome | 0.14 |
| syndromic X-linked intellectual disability | 0.11 |
| amyotrophic lateral sclerosis | 0.04 |
| cold-induced sweating syndrome | 0.01 |
| Cornelia de Lange syndrome | 0.00 |
| multiple congenital anomalies-hypotonia-seizures syndrome | 0.00 |
| congenital disorder of glycosylation type II | 0.00 |
| autosomal dominant non-syndromic intellectual disability | 0.00 |
| Coffin-Siris syndrome | 0.00 |
| coenzyme Q10 deficiency disease | 0.00 |
| congenital disorder of glycosylation | 0.00 |
| mitochondrial DNA depletion syndrome | 0.00 |

**Table S4.** Predictions from skin (UBERON:0002097) model on list of DO terms relevant to the skin.

| DO disease term | Probability |
|---|---|
| oculocutaneous albinism | 0.91 |
| Griscelli syndrome | 0.18 |
| Waardenburg's syndrome | 0.01 |
| tuberous sclerosis | 0.00 |
| dyskeratosis congenita | 0.00 |

### Supplemental Notes

Here we include instances of samples (GSM) and datasets (GSE) wherever their descriptions illustrate points raised in *Discussion*. These instances are grouped into cases where different methods have different performances.

#### 1. Examples where *only* NLP-ML but not TAGGER or MetaSRA correctly annotate samples

- 1.1. GSM139217 - “Cervical carcinoma: None. Age: 66 yrs Gender: Female Ethnicity: Caucasian control_B1, empty vector synthetic_RNA biotin control_B1, empty vector Homo sapiens E-GEOD-5993”
- 1.2. GSM280991 - “HEPG2: Performed by Operator 1 from Site10 at Proficiency stage. HepG2 (liver carcinoma) cell line Proficiency_Site10_21 synthetic_RNA biotin Proficiency_Site10_21 Homo sapiens E-GEOD-11135”

#### 2. Examples where *only* NLP-ML correctly annotate samples but TAGGER, MetaSRA, and expression-based ML do not

- 2.1. GSM267045 - “Genotype:normal well-differentiated primary airway bronchial culture: Gene expression data from human airway epithelium treated as described. Genotype:normal synthetic_RNA well-differentiated primary airway bronchial culture biotin HBE_t0_code1 Homo sapiens E-GEOD-10592”
- 2.2. GSM280991 - “HEPG2: Performed by Operator 1 from Site10 at Proficiency stage. HepG2 (liver carcinoma) cell line Proficiency_Site10_21 synthetic_RNA biotin Proficiency_Site10_21 Homo sapiens E-GEOD-11135”
- 2.3. GSE11881 - "Immunosuppressive drugs can be completely withdrawn in up to 20% of liver transplant recipients, commonly referred to as “operationally” tolerant. Immune characterization of these patients, however, has not been performed in detail, and we lack tests capable of identifying tolerant patients among recipients receiving maintenance immunosuppression. In the current study we have analyzed a variety of biological traits in peripheral blood of operationally tolerant liver recipients in an attempt to define a multiparameter “fingerprint”™ of tolerance. Thus, we have performed peripheral blood gene expression profiling and extensive blood cell immunophenotyping on 16 operationally tolerant liver recipients, 16 recipients requiring on-going immunosuppressive therapy, and 10 healthy individuals. Microarray profiling identified a gene expression signature that could discriminate tolerant recipients from immunosuppression-dependent patients with high accuracy. This signature included genes encoding for T-cell and NK receptors, and for proteins involved in cell proliferation arrest. In addition, tolerant recipients exhibited significantly greater numbers of circulating potentially regulatory T-cell subsets (CD4+CD25+ T-cells and Vd1+ T cells) than either non-tolerant patients or healthy individuals. Our data provide novel mechanistic insight on liver allograft operational tolerance, and constitute a

first step in the search for a non-invasive diagnostic signature capable of predicting tolerance before undergoing drug weaning. Experiment Overall Design: The complete database comprised the expression measurements of 54 675 genes for nine operationally tolerant (TOL) and eight immunosuppression-dependent (ID) samples."

#### 3. Examples where *all* text-based methods (TAGGER, MetaSRA, and NLP-ML) correctly annotate samples

- 3.1. GSM306886 - “tissue: lymph node tumor DLBCL frozen biopsy: frozen biopsy of lymph node tumor Gene expression data from primary DLBCL biopsy. tissue: lymph node tumor synthetic_RNA DLBCL frozen biopsy biotin DLBCL biopsy, sample 2012 Homo sapiens E-GEOD-12195”
- 3.2. GSM190876 - “Non-cancerous renal cortical tissue from nephrectomized kidney with isolated renal cell carcinoma of patient 1, normal control tissue. PKD1 patient kidney synthetic_RNA biotin normal renal cortical tissue_1 normal renal cell carcinoma Homo sapiens E-GEOD-7869”
- 3.3. GSE10780 - “Analysis of 143 completely histologically-normal breast tissues resulted in the identification of a “malignancy risk” gene signature that may serve as a marker of subsequent risk of breast cancer development. Experiment Overall Design: RNA was extracted from microdissected frozen breast tissues for gene array analysis
- 3.4. GSE1145 - “To establish changes in cardiac transcription profiles brought about by heart failure we collected myocardial samples from patients undergoing cardiac transplantation whose failure arises from different etiologies (e.g. idiopathic dilated cardiomyopathy, ischemic cardiomyopathy, alcoholic cardiomyopathy, valvular cardiomyopathy, and hypertrophic cardiomyopathy) and from “normal” organ donors whose hearts cannot be used for transplants. The transcriptional profile of the mRNA in these samples will be measured with gene array technology. Changes in transcriptional profiles can be correlated with the physiologic profile of heart-failure hearts acquired at the time of transplantation. Keywords: other”
- 3.5. GSM1131542 - "srs478673 primary colorectal cancer primary colorectal cancer amc_13-1 homo sapiens amc_13 colon colorectal cancer stage iv colorectal cancer primary tumor extract 1 illumina hiseq 2000 paired cdna transcriptomic rna-seq total rna srx347896 illumina hiseq 2000 (homo sapiens) sequencing assay srr975560_1 srr975560 primary tumor"
- 3.6. GSM882078 - “young vastus lateralis muscle non-exercised y18 20.0 26.697530864197503 homo sapiens male extract 1 genomic dna le 1 cy3 a-geod-13534 array assay norm 20.0 26.697530864197503”

#### 4. Examples where only expression-based ML correctly annotates samples but all text-based methods (TAGGER, MetaSRA, and NLP-ML) do not

- 4.1. GSM318437 - “EFO_0001266 male bipolar disorder 42 amplified RNA total RNA human neurons isolated from postmortem dorsolateral prefrontal cortex 32h 42 biotin neuronal bipolar disorder neuron_rep12 male Homo sapiens neuronal E-GEOD-12679 Stanley Medical Research Institute”
- 4.2. GSM318430 - “EFO_0001265 female schizophrenia 44 amplified RNA total RNA human neurons isolated from postmortem dorsolateral prefrontal cortex 26h 44 biotin neuronal schizophrenia neuron_rep5 female Homo sapiens neuronal E-GEOD-12679 Stanley Medical Research Institute”
- 4.3. GSE19625 - “Coenzyme Q10 (CoQ10) is an obligatory element in the respiratory chain and functions as a potent antioxidant of lipid membranes. More recently, anti-inflammatory effects as well as an impact of CoQ10 on gene expression have been observed. To reveal putative effects of Q10 on LPS-induced gene expression, whole genome expression analysis was performed in the monocytic cell line THP-1. 1129 probe sets have been identified to be significantly up-regulated ($p < 0.05$) in LPS-treated cells when compared to controls. Text mining analysis of the top 50 LPS up-regulated genes revealed a functional connection in the NFκB pathway and confirmed our applied in vitro stimulation model. Moreover, 33 LPS-sensitive genes have been identified to be significantly down-regulated by Q10-treatment between a factor of 1.32 and 1.85. GeneOntology (GO) analysis revealed for the Q10-sensitve genes a primary involvement in protein metabolism, cell proliferation and transcriptional processes. Three genes were either related to NFκB transcription factor activity, cytokinesis or modulation of oxidative stress. In conclusion, our data provide evidence that Q10 down-regulates LPS-inducible genes in the monocytic cell line THP-1. Thus, the previously described effects of Q10 on the reduction of pro-inflammatory mediators might be due to its impact on gene expression. Whole genome expression profiles were analysed from monocytes pre-incubated with ubiquinone (Q10) before subsequent stimulation with LPS. Stimulated (+LPS) and unstimulated (-LPS) monocytes were used as positive and negative controls, respectively. For every experimental group (3 groups in total), three Affymetrix Human Genome U133 Plus 2.0 arrays were used, thus resulting in the analysis of 9 microarrays.”
- 4.4. GSE15935 - “Cyclophilin binding drugs, NIM811 and cyclosporin A (CsA), inhibit the replication of HCV replicon. We investigated the mode of action of these drugs and identified host factors essential for HCV replication in a subgenomic replicon model. Experiment Overall Design: Cultured Huh7 cell were treated with CsA or NIM811 at different concentrations. Cells were harvested after 12, 24 or 48 hours. The extracted mRNA were hybridized on Affymetrix U133 Plus 2 microarrays.”

#### 5. Examples where *all* text-based methods (TAGGER, MetaSRA, and NLP-ML) correctly annotate samples but expression-based ML does not

- 5.1. GSM299098 - “human: Evaluate gene expression profiles after inducing differentiation in cultured interstitial cystitis (IC) and control urothelial cells.. Female, age 47; Disease status: HB (control (stress incontinence, no bladder pain)); Culture medium: KM (proliferating medium) bladder human cultured bladder HB KM, subject 8 47 synthetic_RNA normal female human cultured bladder HB KM, subject 8 proliferation medium patient 8 biotin Homo sapiens urothelial cell E-GEOD-11839”
- 5.2. GSM300205 - “brain, hippocampus, male, 83 years: Brain from cognitively intact individual.. Individual: 28, C; Brain region: hippocampus; Gender: male; Age: 83 years Hippocampus_male_83yrs_indiv28 synthetic_RNA biotin Hippocampus_male_83yrs_indiv28 Homo sapiens E-GEOD-11882”
- 5.3. GSE18696 - “Specific vulnerability of neurons in the human entorhinal cortex has been associated with the onset of disease. Gene expression is analyzed to define the molecular characteristic of those neurons. Experiment Overall Design: Human tissue collection and dissection. Brain samples were collected from four individuals with no clinical evidence of neurological disease and no neuropathological evidence of neurodegeneration. Tissue samples were obtained from the neurological tissue bank (UIPA) and from the Neurological Research Tissue Bank (BTIN, Madrid). The mean postmortem interval (PMI) of the tissue was 6 h and each subject died in hospital due to either cardiac or infectious diseases. The tissue was obtained according to local ethical and legal regulations concerning the use of human post-mortem tissue for biomedical research. Frozen tissue samples were collected from the entorhinal cortex [EC; Brodmann area (BA) 28], at coronal level 27 of the Atlas of Paxinos. Tissue samples corresponded to either upper (CES), lower (CEI) or the entire EC (CET). Two adjacent vertical columns comprising the full thickness of the EC were dissected under magnification with a Leica M50 stereomicroscope. One of them was then divided into two blocks, corresponding to CES and CEI, while the remaining column was processed as CET. RNA sample preparation. Cerebral tissue was homogenised in liquid nitrogen with a pestle and mortar, and the total RNA was isolated using the RNeasy Mini Kit and QIAshredder (Qiagen). The total RNA concentration and purity were determined using an Agilent2100 Bioanalyzer (Agilent Biotechnologies, Palo, Alto, CA) and by agarose gel electrophoresis. Subsequently, cDNA was synthesized using the One-Cycle cDNA Synthesis kit (Affymetrix), according to the protocol described in the Expression Analysis Technical Manual. Biotinylated cRNA probes were generated from each cDNA sample following the IVT Labeling kit instructions (Affymetrix), and the cRNA synthesized was purified with the GeneChip Sample Cleanup Module (Affymetrix). The concentration and purity of the biotinylated cRNA was determined using an Agilent2100 Bioanalyzer (Agilent Biotechnologies, Palo, Alto, CA) and by agarose gel electrophoresis.”
- 5.4. GSE6257 - “The exit of antigen-presenting cells (APC) and lymphocytes from inflamed skin to afferent lymph is vital for the initiation and maintenance of dermal immune responses. How such exit is achieved and how cells transmigrate the distinct endothelium of

lymphatic vessels is however unknown. Here we show that inflammatory cytokines trigger activation of dermal lymphatic endothelial cells (LEC) leading to expression of the key leukocyte adhesion receptors ICAM-1, VCAM-1 and E-selectin, as well as a discrete panel of chemokines and other potential regulators of leukocyte transmigration. Furthermore, we show that both ICAM-1 and VCAM-1 are induced in the dermal lymphatic vessels of mice exposed to skin contact hypersensitivity where they mediate lymph node trafficking of DC via afferent lymphatics. Lastly, we show that TNF_-stimulates both DC adhesion and transmigration of dermal LEC monolayers in vitro and that the process is efficiently inhibited by ICAM-1 and VCAM-1 adhesion-blocking mAbs. These results reveal a CAM-mediated mechanism for recruiting leukocytes to the lymph nodes in inflammation and highlight the process of lymphatic transmigration as a potential new target for anti-inflammatory therapy. Experiment Overall Design: Global gene expression profile of normal dermal lymphatic endothelial cells cultured in media alone (no TNF) compared to that of normal dermal lymphatic endothelial cells stimulated with TNFalpha, 1 ng/ml for 48h.Triplicate biological samples were analyzed from human lymphatic endothelial cells (3 x controls; 3 x TNF treated) and a single sample analyzed from mouse lymphatic endothelial cells (1 x controls; 1 x TNF treated)."

#### 6. Examples where combining expression-based ML and NLP-ML leads to more correct sample annotations compared to either individual method

- 6.1. GSM103559 - "We examined the effects of 48h of knee immobilization on alterations in mRNA and protein in human skeletal muscle.  Biopsies were taken from the vastus lateralis muscle of five men (20.4 + 0.5 years) before and after 48h immobilization. vastus lateralis skeletal muscle knee PHI-CTR-4UP-S2 synthetic_RNA biopsy male PHI-CTR-4UP-S2 subject 4 biotin Homo sapiens E-GEOD-5110 pre-immobilization"
- 6.2. GSM101652 - "Brain tissue from patient NOB1228:  Brain tissue from glioblastopma patient NOB1228. primary  NOB1228 synthetic_RNA biotin NOB1228 Homo sapiens E-GEOD-4536"
- 6.3. GSE13671 - "Female BRCA1 mutation carriers have a nearly 80% probability of developing breast cancer during their life-time. We hypothesized that the breast epithelium at risk in BRCA1 mutation carriers harbors mammary epithelial cells (MECs) with altered proliferation and differentiation properties. Microarray studies revealed that PMEC colonies from BRCA1 mutation carriers anticipate expression profiles found in BRCA1-related tumors, and that the EGFR pathway is upregulated in BRCA1 mutation carriers compared ton non BRCA1 mutation carriers.  Keywords: Class comparison and pathway analysis 10 colonies were collected and RNA was isolated using the Absolutely RNA Nanoprep kit, Stratagene. The arrays included duplicates from four normal controls and from two BRCA1 mutation carriers and single arrays from another two BRCA1 mutation carriers."
- 6.4. GSE25414 - "Genetic factors contribute to the development of ischemic stroke but their identity remains largely unknown. We tested the association with ischemic stroke of 210 single nucleotide polymorphisms (SNPs) associated with pathways functionally related to stroke. We observed an association between the rs7956957 SNP in LRP1 and next

performed microarrays analysis in healthy individuals to investigate possible associations of LRP genotypes with the expression of other genes. Twelve blood samples were obtained from twelve different healthy subjects carrying different genotypes for the rs7956957 SNP of the LRP1 gene (GG, CG or CC). Blood was extracted from 12 subjects. EDTA tubes were centrifuged at 3500rpm for 15 min to obtain the white blood cell fraction and the samples were stored at -80ºC until RNA isolation. RiboPureTM -Blood kit from Ambion (Ambion, Woodward st. Austin, USA) was used to extract total RNA following manufacturer's instructions. Globin RNA from erythrocytes that causes interference was extracted using the Globin-Clear kit (Ambion, Woodward st. Austin, USA)."

- 6.5. GSM456307 - "nan 8076-8077 homo sapiens universal human reference total rna extract 1 total rna le 1 cy5 a-geod-9190 norm not specified not specified not specified not specified not specified not specified not specified"
- 6.6. GSM599874 - "saliva 10-016c cerebral palsy homo sapiens saliva proband male extract 1 genomic dna le 1 cy3 and cy5 a-geod-20641 array assay cerebral palsy proband male"
